# Influence of weather on dragonfly migration and flight behaviour along the Baltic coast

**DOI:** 10.1101/2020.09.03.281253

**Authors:** Aline Knoblauch, Marco Thoma, Myles H. M. Menz

## Abstract

Despite mass movements of dragonflies being documented for decades, the influence of weather on the movement decisions and movement intensity of dragonflies has rarely been studied. Here, we investigate the influence of local weather conditions on flight behaviour of dragonflies in Europe, taking advantage of large movements of dragonflies occurring along the Baltic Sea coast of Latvia. Firstly, we performed orientation tests with individual dragonflies of two commonly captured species, *Aeshna mixta* and *Sympetrum vulgatum*, in order to determine if dragonflies showed directed flight and whether flight direction was independent from wind direction. *Aeshna mixta* displayed a uniform mean southward orientation (166.7°), independent from prevailing wind directions, whereas *S. vulgatum* did not show a uniform orientation. Secondly, we investigated the influence of weather conditions on the abundance of dragonflies captured. Behavioural differences in relation to weather conditions were observed between *A. mixta* and the two smaller *Sympetrum* species (*S. vulgatum* and *S. sanguineum*). Generally, temperature, cloud cover and wind direction were the most important predictors for migration intensity, with temperature positively influencing abundance and cloud cover negatively influencing abundance. *Aeshna mixta* appeared to select favourable tailwinds (northerlies), whereas hourly abundance of *Sympetrum* increased with more easterly winds. Our results provide important information on the influence of local weather conditions on the flight behaviour of dragonflies, as well as evidence of migration for *A. mixta* and most likely some *Sympetrum* species along the Baltic coast.

## INTRODUCTION

Migration has evolved independently in a multitude of taxa within many of the major animal lineages (Alerstam et al. 2003; Dingle and Drake 2007; Bauer and Hoye 2014; Dingle 2014). Insects are the most abundant and diverse group of terrestrial migrants (Holland et al. 2006; Dingle and Drake 2007; Chapman et al. 2015; Hu et al. 2016; Satterfield et al. 2020), yet migratory behaviour has only been well studied in relatively few insect taxa, often restricted to iconic species, such as the Monarch butterfly, *Danaus plexippus* (Flockhart et al. 2013) or agricultural pests (Johnson 1969; Drake and Reynolds 2012; Chapman et al. 2015; Jones et al. 2019) and beneficial species (Wotton et al. 2019; Gao et al. 2020). Over the last 50 years, new tools have been developed to study insect migration at a higher resolution, for example using vertical-looking radars (Chapman et al. 2003, 2011a; Drake and Reynolds 2012), and radio-telemetry (Wikelski et al. 2006; Knight et al. 2019). Intrinsic markers such as stable isotopes have also been used to successfully track population-scale movements across the migratory cycle (Hobson et al. 2012; Flockhart et al. 2013; Stefanescu et al. 2016; Hallworth et al. 2018). These approaches have highlighted the importance of insect migration with respect to biomass movements and ecological impacts (Hu et al. 2016; Wotton et al. 2019) and have provided information regarding migratory routes and specific parameters such as migration height or flight speed (Drake and Reynolds 2012; Chapman et al. 2015). And yet, insect migration is still largely unquantified, and the migratory behaviour and movements of many species remains elusive. Consequently, the use of more traditional methodology, such as flight interception traps and systematic counts can provide critical baseline information for determining population trends, migratory phenology, and behaviour in relation to local weather and topography (Brattström et al. 2008; Krauel et al. 2015).

Many insects are able to select suitable conditions for their migration (Brattström et al. 2008; Chapman et al. 2011b; Drake and Reynolds 2012, Gao et al. 2020) and factors such as temperature and pressure can influence movement behaviour, as well as provide cues for the initiation and termination of migration (Johnson 1969; Wikelski et al. 2006; Brattström et al. 2008; Bauer and Klaassen 2013). While many weak-flying smaller insects, for example aphids, have their migratory displacements dictated mostly by wind (Chapman et al. 2011a,b; Hu et al. 2016; Wainwright et al. 2017; Huestis et al. 2019), larger insects such as moths, butterflies and dragonflies are, to a certain extent, able to control their direction relative to the ground by (partially) compensating for drift as well as exploiting tail-winds (Srygley 2003; Chapman et al. 2008). Some large, low-flying diurnal insects – such as butterflies and dragonflies – often migrate within their ‘flight boundary layer’ – the zone extending up from the ground where the ambient wind speed is lower than the insect’s airspeed (Srygley and Dudley 2008). For these migrants, some proximate weather variables, for example air temperature, wind speed and direction or cloud cover, seem to be recurrent cues which initiate or maintain migratory movements (Wikelski et al. 2006; Brattström et al. 2008; Chapman et al. 2015). Furthermore, aerial migration of some insects often concentrates along topographic elements that may act as barriers or bottlenecks, funnelling migrating animals (Becciu et al. 2019). For example, large concentrations of migrating insects can be observed in areas such as mountain chains, alpine passes (Lack and Lack 1951; Aubert 1962, 1964; Borisov 2009; Thoma and Althaus 2015) or coastlines (Russell et al. 1998; Corbet 1999; Brattström et al. 2008).

While there have been numerous anecdotal reports of dragonfly movement and migration worldwide, beginning in 1494 (Calvert 1893), few studies have systematically documented this phenomenon (Feng et al. 2006; May and Matthews 2008; Shapoval and Buczyński 2012; Hallworth et al. 2018), and relatively little is known about the detailed behaviour and ecology of dragonfly migration (Dumont and Hinnekint 1973; Russell et al. 1998; Corbet 1999; Parr 1996, 2010; May 2013). Corbet (1999), provided a valuable baseline for dragonfly migration research by classifying different types of nontrivial flights. More recently, the use of radio telemetry has provided insights into migration behaviour on an individual level, with information on speed, direction and preferred weather conditions of migrating *Anax junius* in the United States (Wikelski et al. 2006; Knight et al. 2019). In the last decade, effort has been made to document the migratory routes and dynamics of the globally distributed *Pantala flavescens* (Anderson 2009; Chapman et al. 2015) through the use of radar (Feng et al. 2006), stable isotopes (Hobson et al. 2012) and molecular tools (Troast et al. 2016). However, much of our knowledge of dragonfly migration is based on these two iconic species. Dragonflies impact both aquatic and terrestrial ecosystems (Van Buskirk 1988). Their role as natural enemies of pests, such as mosquitoes or aphids, as prey for birds and amphibians (Corbet 1999), but also as bioindicators of freshwater quality (Bulankova 1997, Smith et al. 2007, Da Silva Monteiro Júnior et al. 2015), makes understanding their dynamics and movements important for understanding ecosystem functioning.

By taking advantage of large, annually recurring autumn movements of dragonflies along the Baltic Sea coast, we investigated dragonfly orientation, flight phenology and behaviour, with respect to weather conditions, to provide evidence that the movements observed could be attributed to (long-distance) migration. Specifically, we (1) investigated if dragonflies showed directional flight, and if this is independent from wind direction, (2) investigated how weather conditions influence the strength of dragonfly movement, and (3) explored how movement phenology in relation to weather conditions varied between species.

## MATERIALS AND METHODS

### Study site and sampling

The study took place at the Pape ornithological station in south-western Latvia (56°09’59”N, 21°01’03”E; Figure 1a). The station is situated on the Baltic Sea coast with Pape Lake to the east. Pape Lake is situated parallel to the sea, possibly generating a bottleneck effect on migrating birds and insects (M. Briedis pers. comm.), which often avoid flying over open waters (Alerstam and Christie 1991; Corbet 1999; Becciu et al. 2019), thereby funnelling the migrants along the strip of land between the lake and the sea (von Rintelen 1997; Figure 1a).

**Figure 1.**
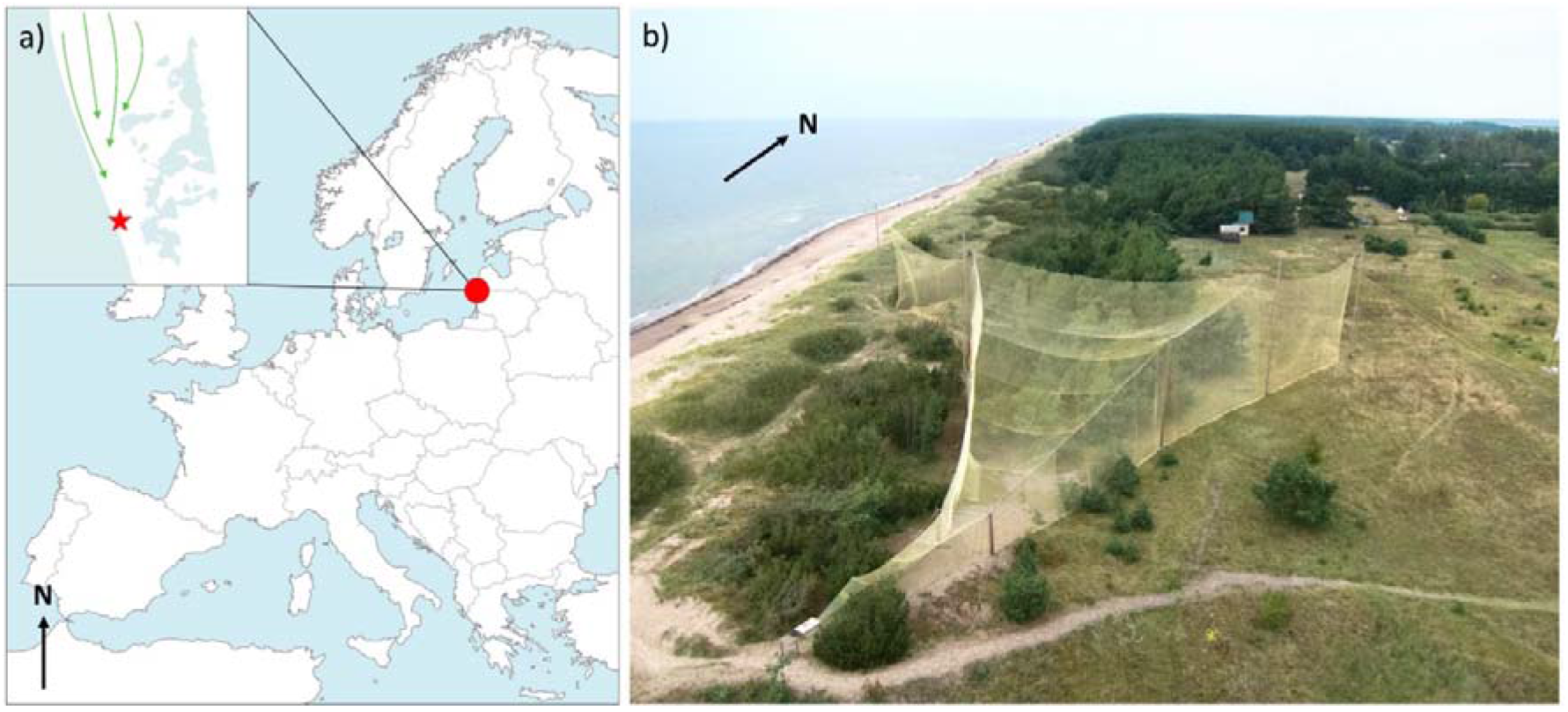
(a) Location of the Pape ornithological research station, Latvia. The red star indicates the position of the Heligoland trap. The green arrows represent the hypothetical flyways due to a bottleneck effect exerted on migrating dragonflies by the Baltic Sea and Pape Lake. (b) Photograph of the large Heligoland trap (Photo: Jasja Dekker).

During summer and autumn, the station successively operates two funnel traps that open to the North (Figure 1b) – also called Heligoland or “Rybachy” traps. These traps are primarily used to catch migrating birds and bats (Šuba et al. 2012; Voigt et al. 2012), but also trap a large number of migrating insects, in particular dragonflies (von Rintelen 1997). The two traps are of different size and are situated at the same location. Dragonflies were caught between 13 August and 9 September 2016 using the large Heligoland trap (entrance size 15 m high x 35 m wide x 40 m long). From 11 September to 9 October, the larger trap was replaced by a smaller trap (entrance size 6 m high x 11.40 m wide x 28 m long). Trap replacement is the standard procedure of the ringing station in order to cope with increase in capture rates of birds over the course of the autumn season. The larger trap is adjustable in height to prevent weather damage to the structure. The height of the trap opening was noted every hour. The traps both have a mesh size of 2 x 2 cm and end in a box (100 x 40 x 40 cm) from which animals can easily be collected. The box was emptied hourly between 08:00 and 18:00 hrs (UTC+2) each day. At 18:00, the rest of the trap was also searched, and the remaining dragonflies were removed and counted. The change from the larger to the smaller trap design caused a sharp reduction of the number of dragonflies captured (Fig. S1). Therefore, we excluded the data using the smaller trap from statistical analyses.

Dragonflies were identified to species level and sexed. Before release, dragonflies were marked on the upper side of the wings using waterproof, coloured paint markers (Edding 4000) with a week-specific colour code to determine the proportion of animals recaptured in the trap, and potentially provide insight into movements through re-sightings of marked individuals. Colour-marking the wings of dragonflies is an effective method for mark-recapture studies and has previously been used for determining local movements and population dynamics (Jacobs 1955; Borisov 2009; Keller et al. 2010; Kharitonov and Popova 2011). The putative migratory status of the dragonfly species recorded was based on Corbet (1999), Jović et al. (2010) and Schröter (2011) (Table 1).

**Table 1.**
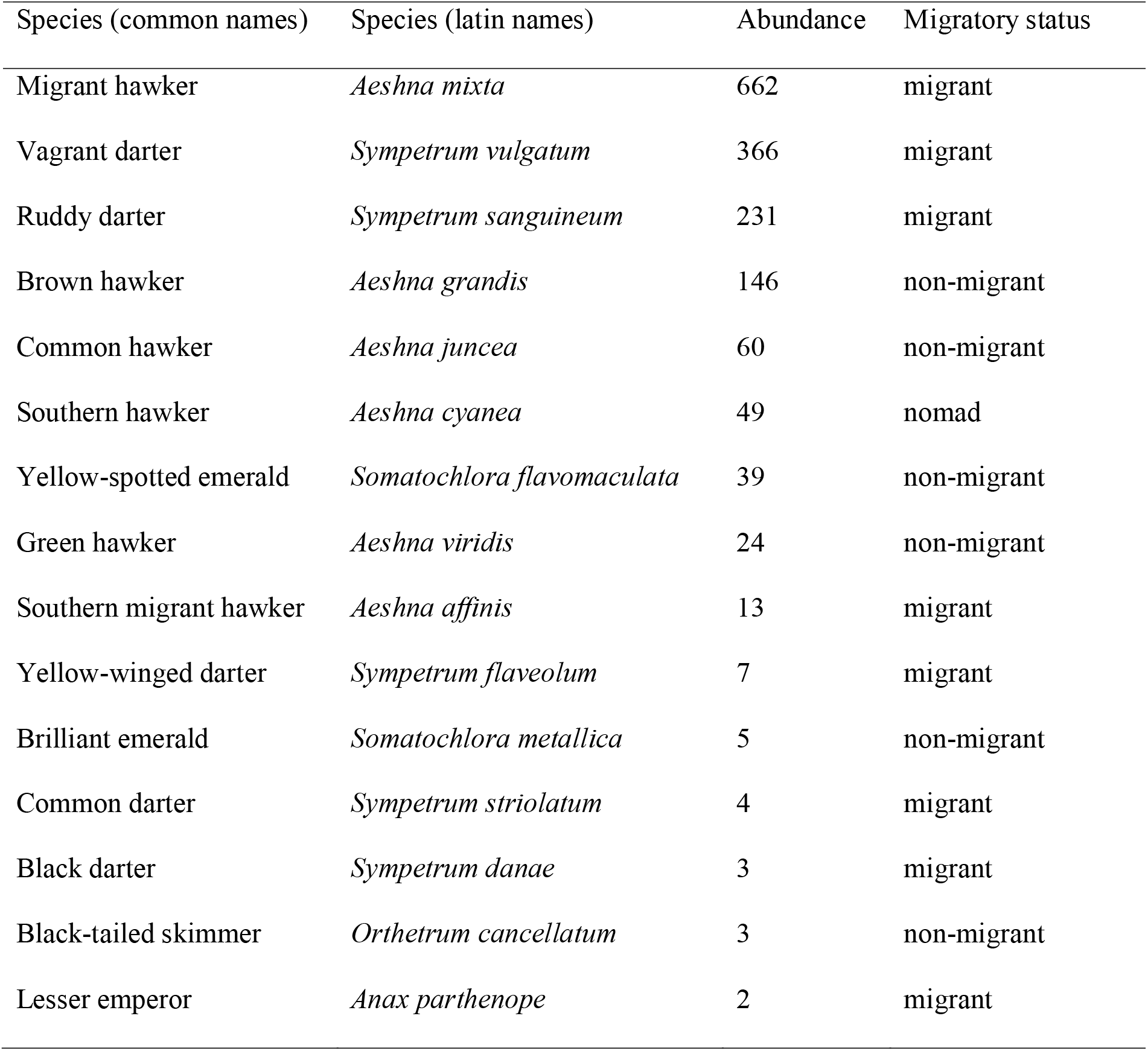
Abundance and migratory status of dragonflies captured at Pape, Latvia. Migrant as a status encompasses obligate, partial and facultative strategies, as in some cases status is not clear and prone to change depending on latitude, for example. Non-migrant species are species for which no migration data has been found in the literature (Corbet, 1999; Jović et al., 2010; Schröter, 2011). Data are from the box at the end of the larger trap (see Methods).

### Orientation tests

Flight orientation tests were carried out in the middle of an open field adjacent to the trap. The experimental site was located at least 40 m from the nearest visual cues (trees or bushes). We used a circular flight arena made of black mesh (38 cm in diameter and 30 cm high), placed 45 cm above the ground. A video camera (Sony Handycam DCR-SR200, 4 MP, 60 fps) was positioned above the arena to record flight behaviour. Experiments were conducted with three of the most commonly recorded species, *Aeshna mixta, Aeshna grandis* and *Sympetrum vulgatum*. Both *A. mixta* and *S. vulgatum* are considered to be potential migrants (Dyatlova and Kalkman 2008; Samraoui et al. 2012; Popova and Haritonov 2014) and were captured in high numbers during the season (Table 1).

Dragonflies were released individually into the flight arena immediately after capture and filmed for 15 min. The flight arena was rotated 90° between each experiment in order to randomise potential effects of the arena on flight direction. The videos were analysed using Solomon Coder software (version beta 16.06.26, Péter 2011). The body orientation of the dragonfly was recorded for each video frame and estimated to the nearest 45°.

#### Weather data

Hourly weather data (air temperature [°C], wind speed [km/h], wind direction [°], air pressure [bar] and humidity [%]) was obtained from a weather station (Davis Vantage Pro 2) situated 15 m above the ground, approximately 60 m from the trap. Additionally, cloud cover was recorded every hour on a 0–8 scale, 0 corresponding to no cloud and 8 corresponding to a completely overcast sky.

### Statistical analysis

#### Orientation tests

Analysis of the orientation tests was conducted using the software Oriana 4 (Kovach 1994). As the marking did not have any significant effect on flight direction (Hotelling’s test: *F* = 0, *P* < 0.001), the data of the marked and unmarked dragonflies was pooled for further analyses. Uniformity of the distributions was first tested using a Rayleigh test (Ruxton 2017). We then performed a modified version of the Rayleigh test, also called a *F*-test, to test for uniformity against a unimodal specified mean direction (*μ*). Mean directions were set at 180°, as we hypothesised this to be the main direction during Autumn migration, and 160° which is the direction of the coast at the research station. Sexes were pooled for the analyses. *Aeshna grandis* was excluded from the analysis due to the relatively low number of individuals tested (*N* = 5). For dragonfly species where more than 10 individuals were tested (*A. mixta* and *S. vulgatum*), multi-sample tests were performed to compare their orientation with overhead wind direction during the experiments. These analyses were carried out using a paired *χ*^2^-test in which the classes with zero observations were dropped, more than 20% of the classes having expected frequencies less than five (Kovach 2011). For this analysis, wind direction was converted into the direction towards which the wind blows to match the dragonfly flight direction.

#### The effect of weather on capture rate

All analyses were conducted in R (version 3.3.1., R Core Team 2019) unless otherwise specified. Initially, we performed a Principal Components Analysis (PCA) to investigate whether the dragonfly species clustered based on their phenology. The PCA revealed that the four most common species (*Aeshna grandis, Aeshna mixta, Sympetrum sanguineum, Sympetrum vulgatum*) were separated into two clear clusters, with *A. mixta* and *A. grandis* forming one group, and *S. sanguineum* and *S. vulgatum* forming another (Fig S2). Therefore, subsequent analyses of the influence of weather on abundance were performed based on this grouping. However, there being no mention of *A. grandis* showing migratory behaviour in the available literature, we excluded it from our models.

We used generalized linear mixed effects models in the package “lme4” (Bates et al. 2015) to assess the influence of weather conditions (air temperature [°C], wind speed [km/h], wind direction [°], air pressure [bar], humidity [%] and cloud cover) on the number of dragonflies captured per hour. As wind direction is circular, ranging from 0-360°, for better suitability for linear analysis, the sine and cosine of wind direction were calculated, with the sine corresponding to east-west and cosine corresponding to north-south (Brattström et al. 2008). Trap height and number of days since the start of the sampling were also included in the models. Day number (expressed as ordinal date) was included as a random factor in the models to account for multiple samples per day. Continuous explanatory variables were scaled for the modelling by subtracting the mean and dividing by the standard deviation, using the scale function in R. Models were fitted assuming a Poisson error distribution and checked for overdispersion using the package DHARMa (Hartig 2020). If overdispersion was present (>1.4), we included an observation-level random effect (OLRE) in the model (Harrison 2014). Models were compared using ANOVA and the OLRE was retained if it significantly improved the model fit.

Generalised linear models were used to investigate the influence of weather variables on daily captures, including the set of predictors as for the hourly models described above, as well as day since the beginning of the study. For each weather variable, we calculated the daily mean, between 08:00 and 18:00. We also included the lowest temperature and mean pressure registered on the preceding night (between 18:00 and 06:00 hours). As above, models were initially fitted assuming a Poisson error distribution and tested for overdispersion. If overdispersion was present, we applied a negative binomial model.

For all models, only dragonflies captured in the box at the end of the trap were used for the analysis. Significance of the variables was determined by excluding the variable of interest and comparing the models with and without the variable, using ANOVA. Model fit was visually checked using quantile plots of the residuals. Correlation between the variables used in the models was tested using Pearson correlation tests, with a correlation coefficient threshold of 0.7. No variables were significantly correlated.

## RESULTS

### Orientation tests

The Rayleigh test rejected uniformity of distribution for *A. mixta (Z* = 8.525, *P* < 0.001). Mean flight orientation for the species was 166.67° ± 12.23 (SE) and was significantly different from overhead wind direction during the experiments (χ^2^ = 27.259, *P* < 0.001, Table 3, Figure 2). *Sympetrum vulgatum* showed a mean flight orientation of 205.36° ± 29.88. Uniformity of distribution could not be rejected based on the Rayleigh test, but flight orientation during the tests was significantly different to overhead wind direction (χ^2^ = 19.2, *P* = 0.008, Table 3, Figure 2).

**Figure 2.**
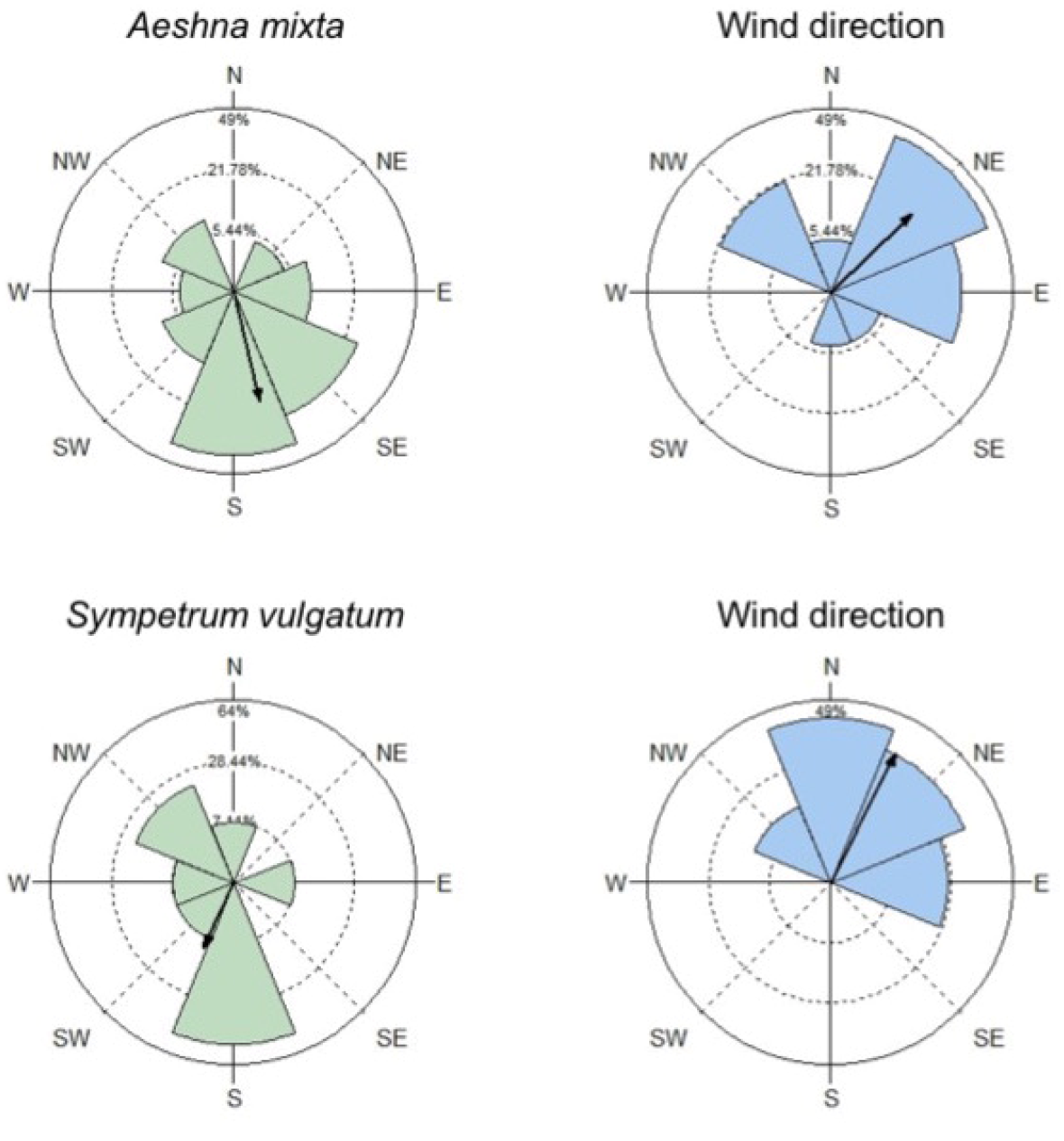
Results of the flight orientation tests. The left column (green) represents the orientation direction of the dragonflies (*A. mixta, N* = 22; *S. Vulgatum, N* = 14). The right column (blue) represents the direction of the wind during these tests. The black arrow represents the mean direction and the wedges the percentage of individuals having their mean direction included in a given wedge. Note that the mean orientation direction for *S. vulgatum* was not significant.

**Table 2.** Results of the flight orientation experiments for *Aeshna mixta* and *Sympetrum vulgatum*. Uniformity of direction was tested using Rayleigh tests. *V*-tests were used to test for uniformity of direction against a unimodal specified mean direction (*μ*) of 180°.

|  | *A. mixta* | *S. vulgatum* |
| --- | --- | --- |
| *N* | 23 | 14 |
| Mean vector ($\mu$) | 166.67° | 205.36° |
| Length of $\mu$ | 0.609 | 0.389 |
| SE of mean | 12.32° | 29.88° |
| *One sample tests* |  |  |
| Rayleigh test (*Z*) | 8.525 | 2.119 |
| Rayleigh test (*P*) | < 0.001 | 0.12 |
| *Multi-sample tests (matched with corresponding wind direction)* |  |  |
| $\chi^2$ pairwise | 27.259 | 19.2 |
| $\chi^2$ pairwise (*P*) | < 0.001 | 0.008 |

**Table 3.** Results of the generalised linear models per dragonfly genus. a) Hourly abundance in relation to weather. b) Daily abundance in relation to weather.

| | | Estimate | SE | df | $\chi^2$ | P value |
| --- | --- | --- | --- | --- | --- | --- |
| **b) *Aeshna*** | Intercept | -16.697 | 2.739 |  |  |  |
|  | Temperature | 1.093 | 0.145 | 1 | 30.550 | <0.001 |
|  | Cloud cover | -0.171 | 0.073 | 1 | 4.615 | 0.032 |
|  | Wind direction (cos) | 0.758 | 0.392 | 1 | 3.930 | 0.047 |
| *Sympetrum* | Intercept | 0.766 | 1.217 |  |  |  |
|  | Cloud cover | -0.396 | 0.092 | 1 | 10.142 | 0.001 |
|  | Trap height | 3.997 | 1.175 | 1 | 8.357 | 0.004 |
| **a) *Aeshna*** | Intercept | -18.831 | 3.659 |  |  |  |
|  | Temperature | 0.814 | 0.128 | 1 | 30.948 | <0.001 |
|  | Cloud cover | -0.117 | 0.050 | 1 | 5.293 | 0.021 |
|  | Wind direction (cos) | 0.490 | 0.161 | 1 | 9.307 | 0.002 |
|  | Humidity | 0.046 | 0.021 | 1 | 4.666 | 0.031 |
| *Sympetrum* | Intercept | -17.844 | 2.991 |  |  |  |
|  | Temperature | 0.947 | 0.155 | 1 | 52.657 | <0.001 |
|  | Cloud cover | -0.165 | 0.075 | 1 | 4.692 | 0.030 |
|  | Wind direction (sin) | 0.581 | 0.226 | 1 | 6.525 | 0.011 |

Significance of variables was determined based on likelihood-ratio tests using ANOVA.
Only significant or variables ( $P < 0.05$ ) were retained in the final models presented here.
Models investigating hourly abundance were fitted with a Poisson error distribution and
included day number as a random factor. An observation-level random factor was included to account for overdispersion. Models investigating daily abundance were fitted with negative binomial models. Model estimates are based on unscaled variables.

### Phenology

In total, 1630 dragonflies were captured in the larger trap (1614 in the end box), of which 75.42% were males and 24.58% females. Overall, 15 species were captured, of which nine are considered to be migratory (Table 1). *Aeshna mixta* accounted for 39.7% of the individuals, followed by *Sympetrum vulgatum* (22.4%), *Sympetrum sanguineum* (14.4%) and *Aeshna grandis* (9.3%). The other species each accounted for less than 4% of the total dragonflies captured. It is to be noted that no damselflies (Zygoptera) were recorded within the trap box.

Out of 2055 individuals marked during the study (147 individuals were caught in the small trap and 278 individuals were caught opportunistically in the surrounds and marked), only 66 (3.2%) were recaptured in the trap (Table S1), with a single individual being reported elsewhere (2 km south of the trap; B. Gliwa, pers. communication). Most of the recaptures (60.6%) took place in the same week the marking was done and the longest time interval between marking and recapture was approximatively one month.

Phenology differed between the four most commonly captured species. The median date of passage for *A. mixta* (26 August, IQR = 21 August – 28 August) and *A. grandis* (25 August, IQR = 22 August – 31 August) tended to be slightly earlier in the season than for the smaller *S. vulgatum* (1 September, IQR = 26 August – 8 September) and *S. sanguineum* (28 August, IQR = 26 August – 1 September, Figure S3). Daily mean capture time was earlier for *A. mixta* (13:07 ± 02:51 SD) and *A. grandis* (12:41 ± 02:06 SD) compared to *S. vulgatum* (14:36 ± 02:00 SD) and *S. sanguineum* (14:44 ± 02:07 SD) (Figure S4).

### The effect of weather on capture abundance

Daily abundance of *A. mixta* increased with temperature (χ^2^ = 30.550, df = 1, *P* < 0.001) and more northerly winds (cosine of wind direction, χ = 3.930, df = 1, *P* = 0.047), and decreased with cloud cover (χ^2^ = 4.615, df = 1, *P* = 0.032, *N* = 25 days, Table 3a, Figure 3). Similarly, hourly abundance of *A. mixta* was positively influenced by temperature (χ^2^ = 30.948, df = 1, *P* < 0.001) and northerly winds (χ^2^ = 9.307, df = 1, *P* = 0.002), and negatively by cloud cover (χ^2^ = 5.293, df = 1, *P* = 0.021), but was also positively influenced by humidity (χ^2^ = 4.666, df = 1, *P* = 0.031, *N* = 251 hours, Table 3b, Figure 4).

**Figure 3.**
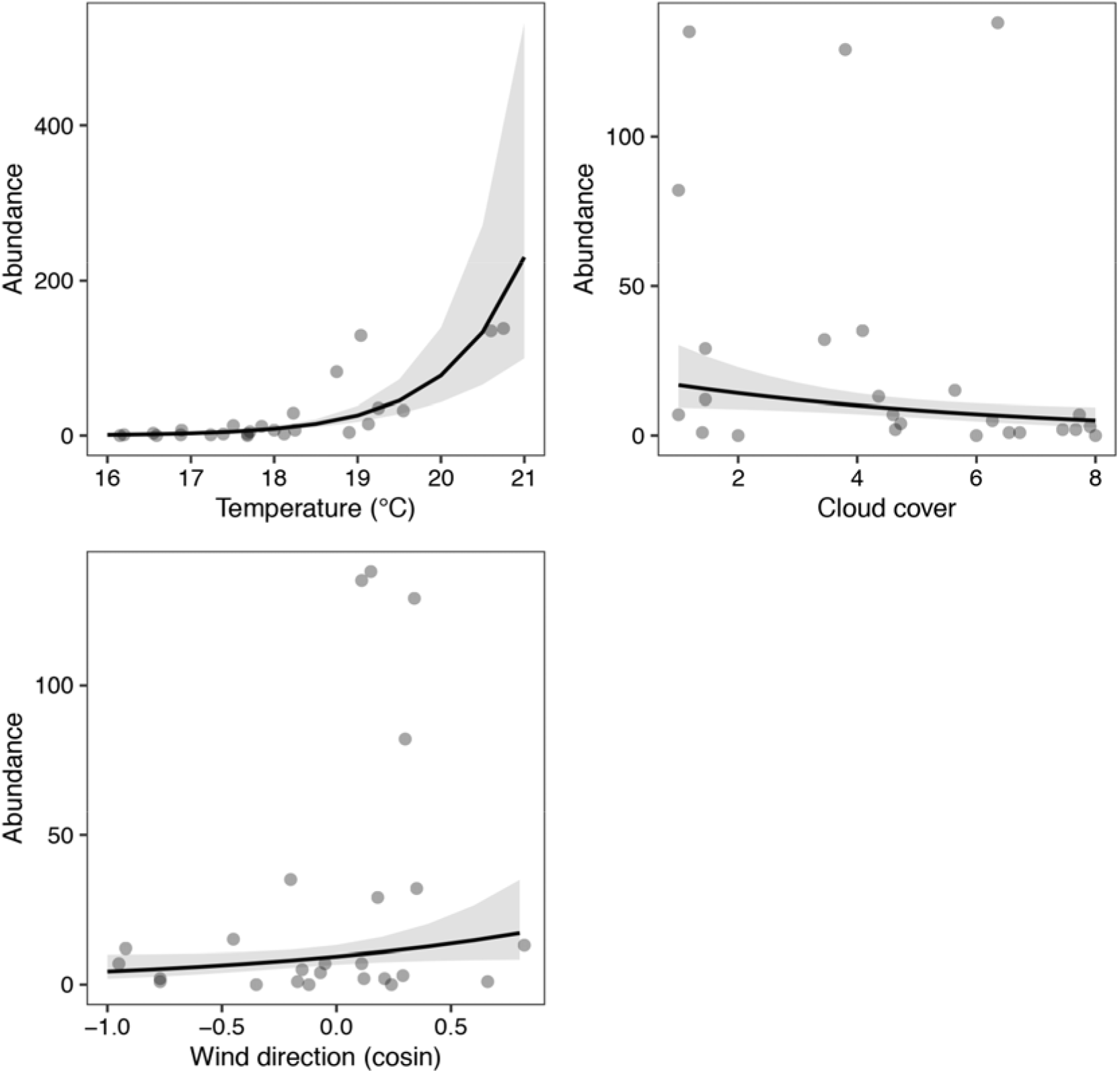
Relationship between weather variables and the daily abundance of *Aeshna mixta*. Results are based on negative binomial regression. The shaded areas represent 95% confidence intervals. For the cosine component of wind direction, (−1) equals south and (+1) equals north.

**Figure 4.**
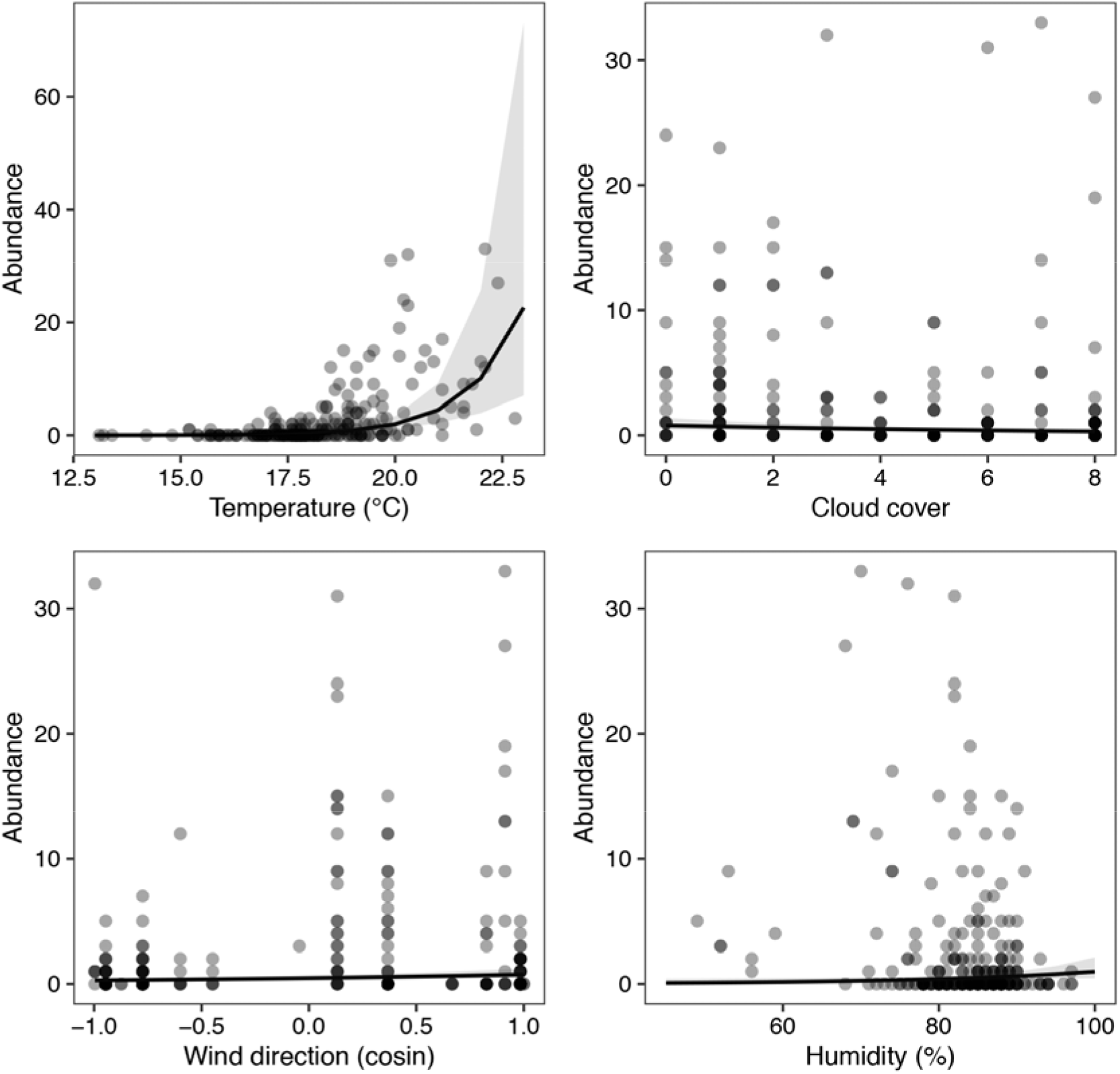
Relationship between the hourly abundance of *Aeshna mixta*, and weather variables. Results are based on a generalised linear mixed model with a Poisson distribution. The shaded areas represent 95% confidence intervals. For the cosine component of wind direction, (−1) equals south and (+1) equals north.

Daily abundance of *Sympetrum* spp. (*S. vulgatum* and *S. sanguineum)* was negatively influenced by cloud cover (χ^2^ = 10.142, df = 1, *P* = 0.001) and positively by trap height (χ^2^ = 8.357, df = 1, *P* = 0.004), with more individuals captured when the trap was fully open (*N* = 25 days, Table 3a, Figure 5). Hourly abundance of *Sympetrum* spp. was positively influenced by temperature (χ^2^ = 52.657, df = 1, *P* < 0.001) and negatively by cloud cover (χ^2^ = 4.692, df = 1, *P* = 0.030, *N* = 251 hours, Table 3b, Figure 6). In contrast to *A. mixta*, hourly abundance of *Sympetrum* spp. increased with more easterly winds (sine of wind direction, χ^2^ = 6.525, df = 1, *P* = 0.011).

**Figure 5.**
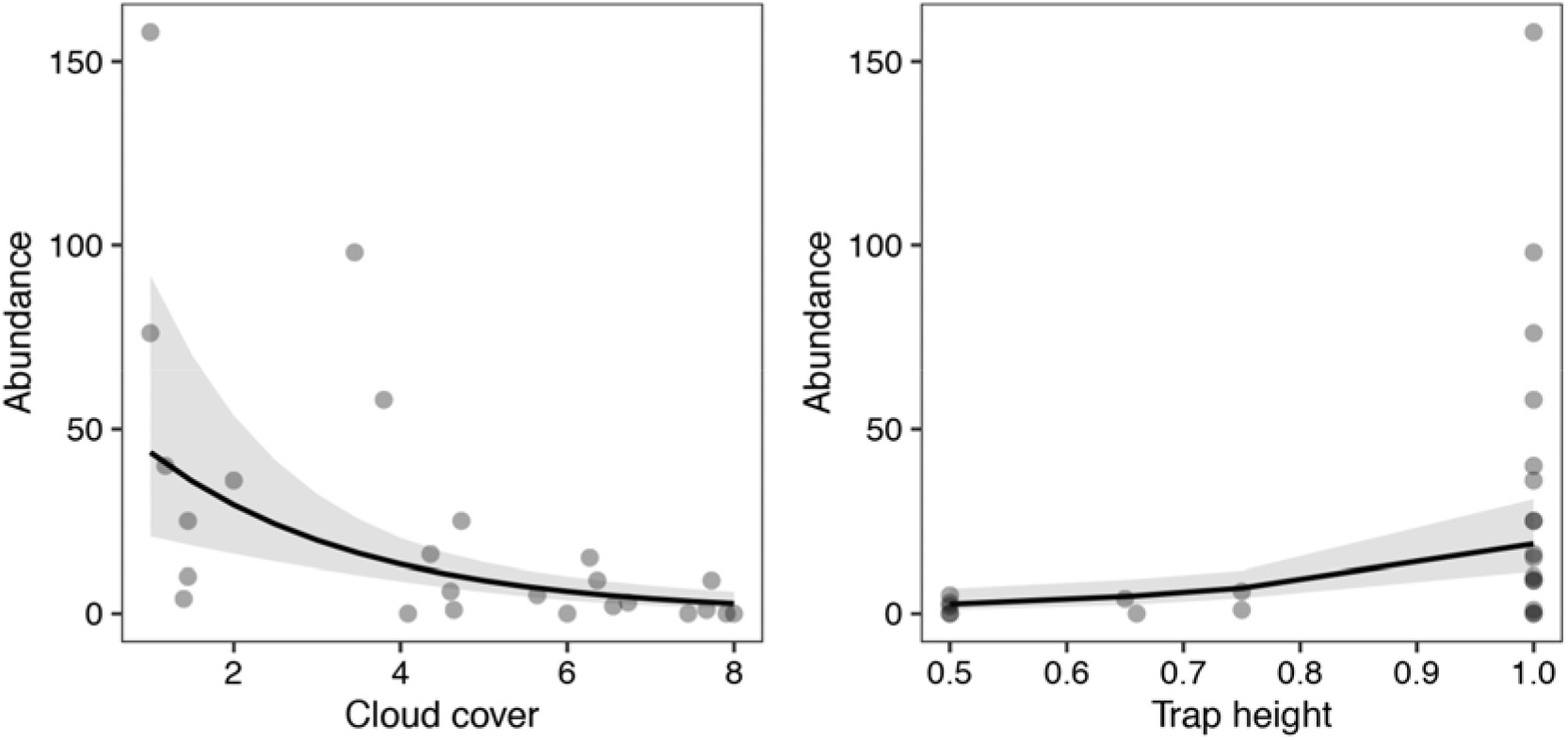
Relationship between weather variables and the daily abundance of *Sympetrum vulgatum* and *Sympetrum sanguineum*. Results are based on negative binomial regression. The shaded areas represent 95% confidence intervals.

**Figure 6.**
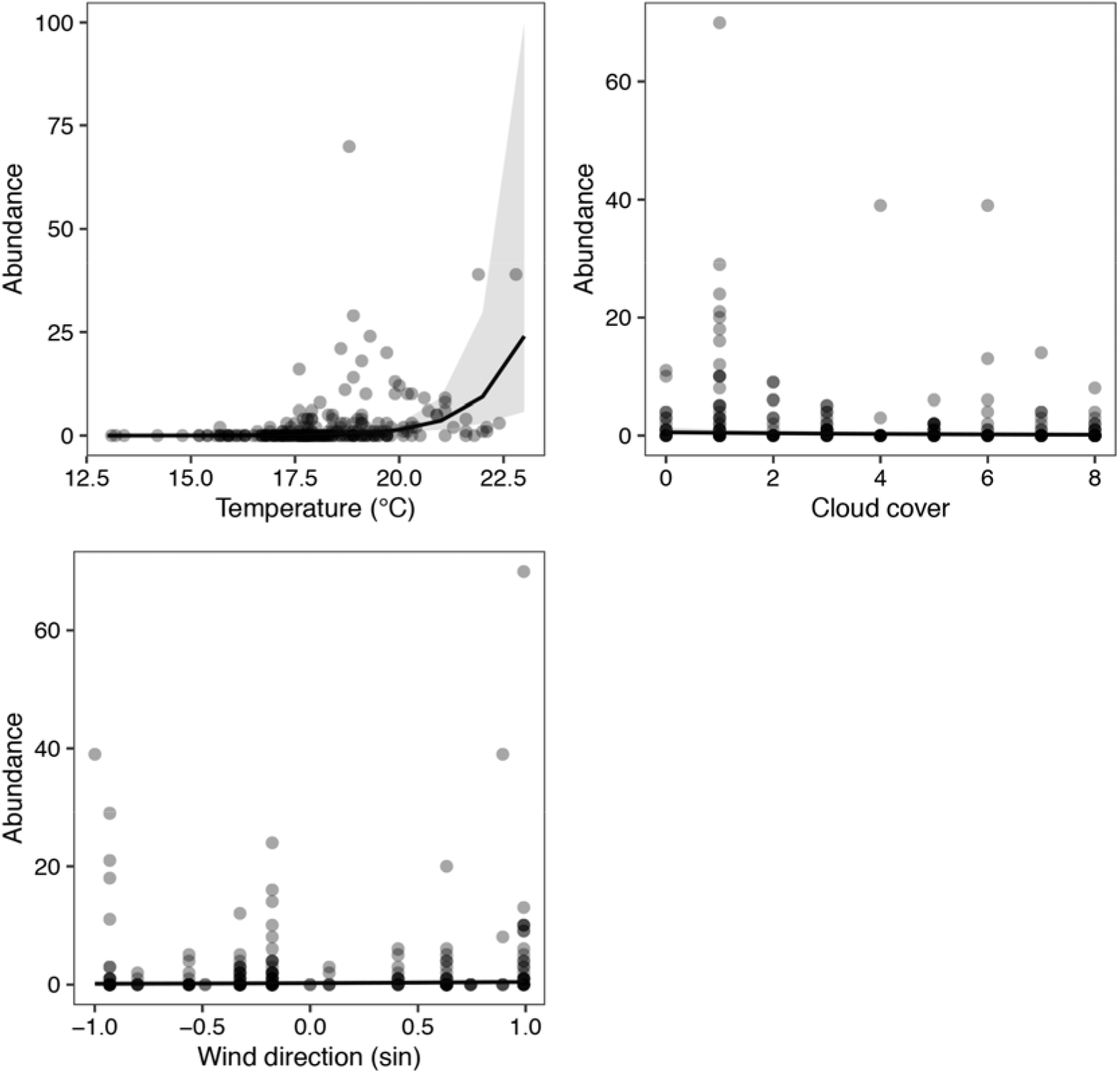
Relationship between the hourly abundance of *Sympetrum* (*Sympetrum vulgatum* and *Sympetrum sanguineum*) and weather variables. Results are based on a generalised linear mixed model with a Poisson distribution. The shaded areas represent 95% confidence intervals. For the sine component of wind direction, (−1) equals west and (+1) equals east.

## DISCUSSION

There is a growing body of evidence for seasonally directed autumn migratory movements in large and medium-sized insects, including day-flying species (Srygley and Dudley 2008; Chapman et al. 2015; Hu et al. 2016; Knight et al. 2019; Wotton et al. 2019; Gao et al. 2020). In dragonflies, directional autumn movements towards the south have been shown for long distance migrant species such as *Anax junius* (Wikelski et al. 2006; Knight et al. 2019). However, to date, relatively few systematic studies on migratory behaviour in dragonflies have been made in Europe (eg. Shapoval and Buczyński 2012). Our orientation tests revealed a strong unimodal flight direction of *Aeshna mixta* towards the south-southeast (167°) in autumn, independent from overhead wind direction. Many insects are known to follow geographical cues on migration (Becciu et al., 2019), and the deviation from true south (180°) observed in *A. mixta’s* orientation could be due to animals following the coastline (160° at Pape). We did not find a significant uniform direction for *Sympetrum vulgatum*. Nonetheless, the mean direction of *S. vulgatum* indicated an orientation towards the south-southwest (205°) and differed from the prevailing overhead winds.

To date, the relationship between dragonfly movement decisions and weather factors has rarely been studied (but see Feng et al. 2006; Wikelski et al. 2016; Knight et al. 2019). We show that temperature and cloud cover were the most important variables affecting hourly and daily abundance of *Aeshna mixta* and the two *Sympetrum* species tested. Both, temperature and cloud cover are well-known predictors for insect migration rates (eg. Brattström et al., 2008), and temperature has been shown to influence groundspeed of migrating *Anax junius* (Knight et al. 2019). This general pattern regarding the influence of temperature and cloud cover is likely to be due to the effect of increased sunshine on flight activity in day-flying insects (Becciu et al. 2019).

Hourly and daily abundance of *A. mixta* was higher in tailwind conditions (in this case, northerlies). Given the predominant southerly flight direction of *A. mixta* revealed in our orientation tests, this pattern might be indicative of the ability to select favourable tailwinds, which would facilitate migration. Many migratory insects such as dragonflies, moths and hoverflies, have been shown to select for favourable winds, possibly as an adaptation to maximize distance covered, optimize trajectories and reduce energy costs in their displacements (Anderson 2009; Alerstam et al., 2011; Becciu et al. 2019; Knight et al. 2019; Gao et al. 2020). Across the study period, the prevailing winds from the SW and W – coming from the Baltic sea – were the strongest and most frequent, while the less common winds coming from the east were of lower speed. The hourly, but not daily abundance of *Sympetrum* increased with easterlies, which may indicate an adaptive response, flying lower to minimize drag and consequently drift due to a reduction in side winds, or to avoid the possibility of being blown out to sea.

While many studies mention that dragonfly migration takes place predominantly in the flight boundary layer (Srygley and Dudley 2008; Chapman et al. 2011a), high-altitude movements up to 1000 m above ground level have been documented for the migratory *Pantala flavescens* (Feng et al. 2006). We therefore cannot be sure whether the increase of passage intensity of *A. mixta* during tail-wind conditions (assuming a general southerly flight direction, based on the results of orientation tests) is limited to ground level or whether passage also intensifies at higher altitudes. In any case, at ringing stations where birds are captured during active migration, songbirds are usually captured in higher numbers when they are facing headwinds (Komenda-Zehnder et al. 2010), flying lower in order to minimize the effect of the wind, which is also the case at Pape (M. Briedis, pers. comm.). Birds tend to fly higher in tailwind conditions in order to take advantage of directionally beneficial winds (Komenda-Zehnder et al. 2010; Becciu et al. 2019). Insects that are strong fliers, such as dragonflies, also use similar mechanisms (Chapman et al. 2015; Becciu et al. 2019). Change in the trap’s height only affected the daily abundance of *Sympetrum* and did not seem to influence the abundance of *A. mixta*. Trap height affecting the capture of *Sympetrum* possibly indicates that either these species fly in a more defined altitudinal layer than the *Aeshna* species, or that they may generally fly higher above the ground. However, our results indicate that *A. mixta* may migrate relatively close to the ground, even in favourable tail-wind conditions, but this warrants further investigation.

The differences observed in the abundance of *Aeshna* and *Sympetrum* throughout the day may be explained by physiological differences found between larger and smaller dragonfly species. The minimum temperature at which flight is possible has been shown to be positively correlated with body weight in dragonflies (May 1976). Thus, for the smaller *Sympetrum* species, the minimum required temperature for flight is higher and hence later in the day.

While the dragonflies captured in the trap may have been a mixture of migrating and local individuals, our results show that there were some differences in the way weather impacts flight behaviour of larger versus smaller dragonfly species. Generally, migration intensity increased with temperature and decreased with cloud cover. Moreover, based on the orientation tests, we have strong support that migration of *A. mixta* and some *Sympetrum* species does take place along the Baltic coast (Shapoval and Buczyński 2012).

Investigation of migration behaviour at altitude, and the origin of individuals – which would provide further insight into migration routes and migratory behaviour – could be answered using a combination of techniques, such as radar (Chapman, Drake, et al., 2011a; Drake & Reynolds, 2012), radio-telemetry (Wikelski et al., 2006, Knight et al. 2019), and stable isotope analysis of wing chitin (Hobson, Soto, Paulson, Wassenaar, & Matthews, 2012; Hallworth et al., 2018).

## Acknowledgements

We thank Oskars Keišs and the Laboratory of Ornithology, Institute of Biology, of the University of Riga for permission to work at the site and assistance with organisation of the fieldwork, Gunārs Petersons for the weather data, and Christian Voigt for bringing the massive dragonfly occurrence at Pape to our attention. Viesturs Vintulis and Mārtiņš Kalniņš provided information on the status of dragonflies in Latvia. We thank D. Račkauskaite and Z. Gasiūnaitė for reporting the observation of a marked specimen. Special thanks to Donāts Spalis for technical support in the field; Lisa Fisler, Martins Briedis, and the Latvian and German volunteers and scientists working at Pape for support during the field study; Wolfgang Nentwig for supporting the study; and Don Reynolds for constructive comments on an earlier version of the manuscript. Fieldwork for this study was conducted under permit Nr.14/2016-E.

