## Supplementary material for "Influence of weather on dragonfly migration and flight behaviour along the Baltic coast"

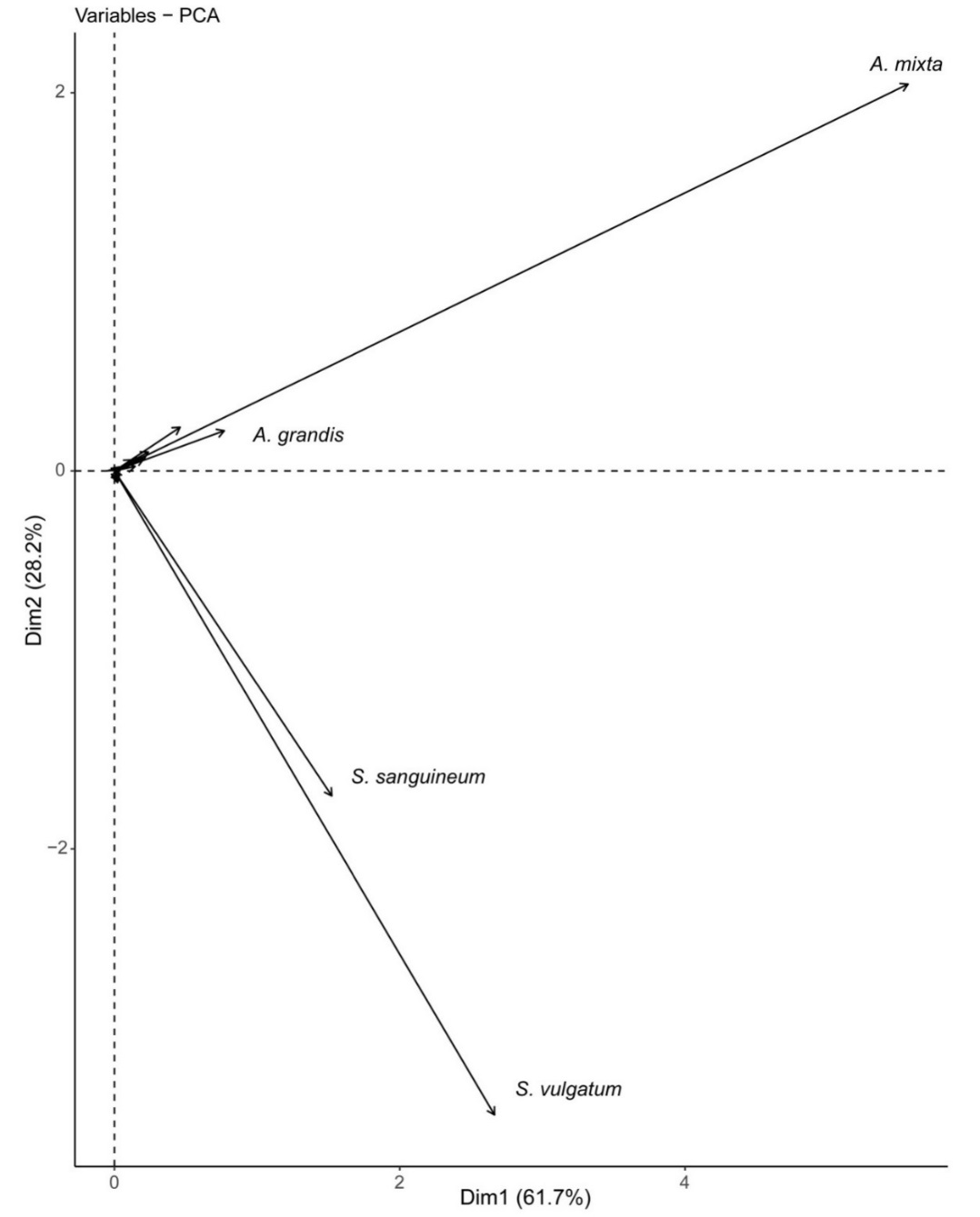

**Figure S1**

Principal Components Analysis of overall migration phenology. The first principal component (Dim1) explains 61.7% of the variance and 89.9% of the variance is accounted for in total by the two first components (Dim1 and Dim2). For better visibility only the four most commonly captured dragonfly species are labelled in this figure.

| **Species** | **Number of recaptures** |
| --- | --- |
| *Aeshna mixta* | 30 |
| *Sympetrum vulgatum* | 12 |
| *Aeshna grandis* | 12 |
| *Sympetrum sanguineum* | 4 |
| *Aeshna cyanea* | 2 |
| *Aeshna juncea* | 2 |
| *Aeshna sp.* | 2 |
| *Aeshna viridis* | 1 |

**Figure S2**

Number of recaptured marked dragonflies per species.

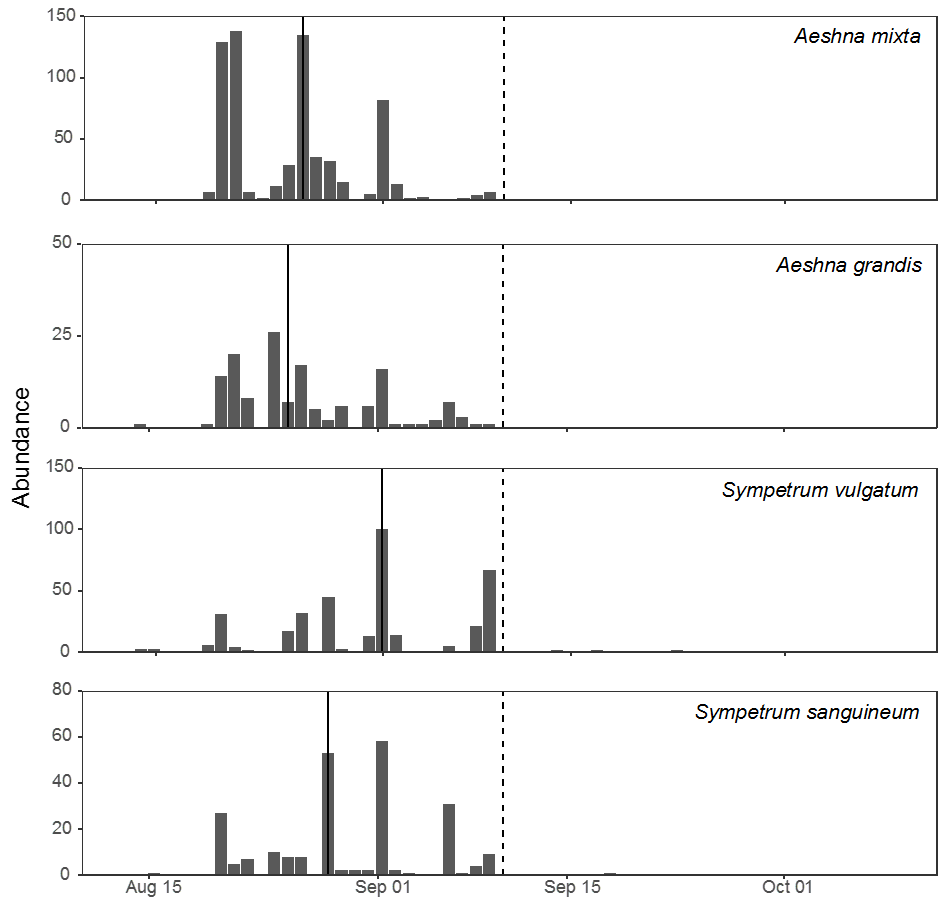

**Figure S3**

Phenology of dragonflies captured at Pape, Latvia during the study period (13 August to 9 October). The vertical dark line represents the median day of passage and the dotted line represents the date at which the large trap was taken down and replaced by the smaller trap.

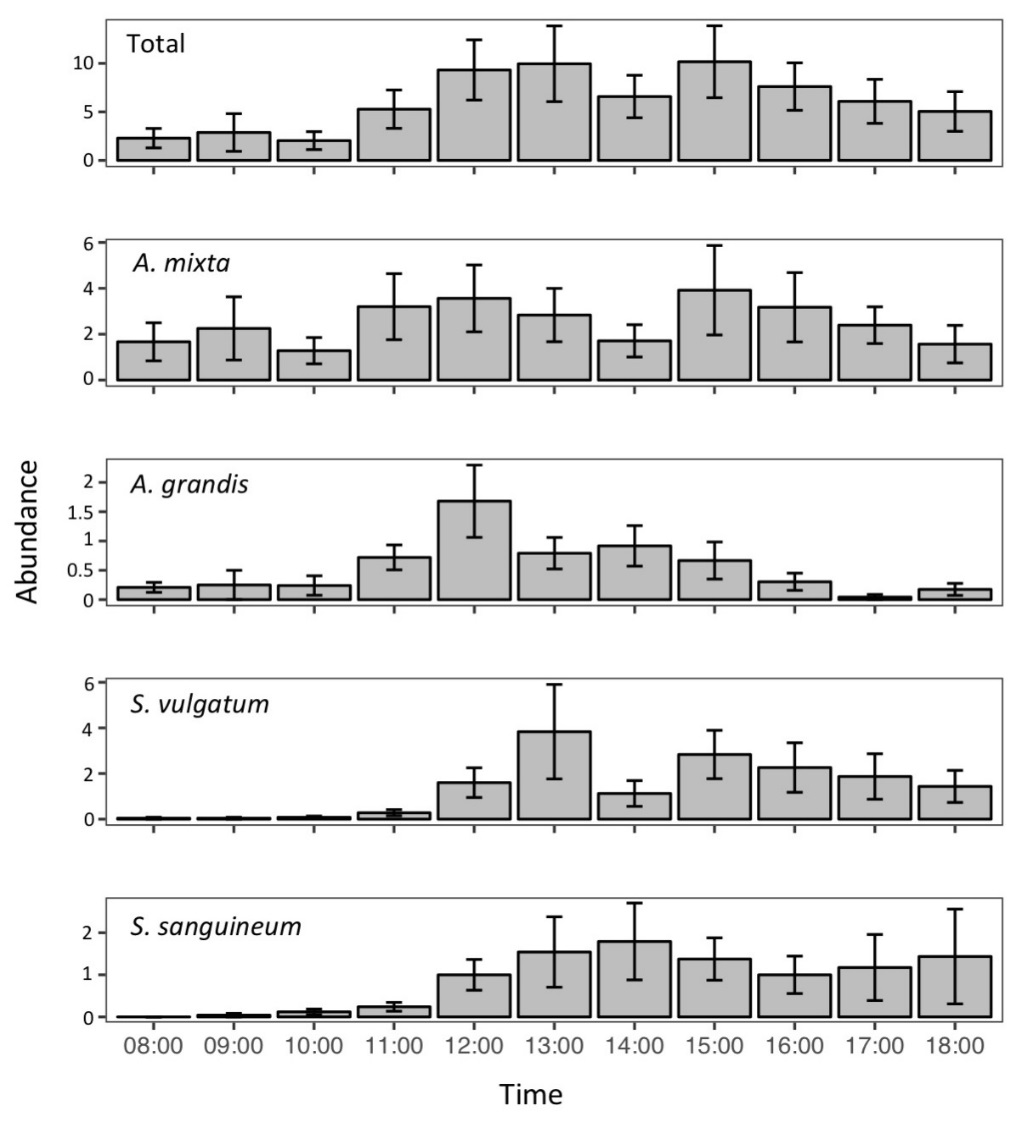

**Figure S4**

Daily phenology from the large trap (13 August to 9 September) of all captured species (Total) and of the four most commonly captured species. The y-axis represents the mean number of individuals captured while the x-axis represents the time of the day at which the trap was emptied. The error bars represent the standard error of the means.
